# Correcting Non-Uniform Milling in FIB-SEM Images with Unsupervised Cross-Plane Image-to-Image Translation

**DOI:** 10.1101/2025.09.29.679411

**Authors:** Yicong Li, Yuri Kreinin, Siyu Huang, Erik W. Schomburg, Dmitri B. Chklovskii, Hanspeter Pfister, Jingpeng Wu

## Abstract

**Motivation:** Focused Ion Beam Scanning Electron Microscopy (FIB-SEM) is an advanced Volume Electron Microscopy technology with growing applications, featuring thinner sectioning compared to other Volume Electron Microscopes. Such axial resolution is crucial for accurate segmentation and reconstruction of fine structures in biological tissues. However, in reality, the milling thickness is not always uniform across the sample surface, resulting in the axial plane looking distorted. Existing image processing approaches often: (i) assume constant section thickness; (ii) consist of multiple separate processing steps (i.e., not in an end-to-end fashion); (iii) require ground truth images for modeling, which may entail significant labor and be unsuitable for rapid analysis.

**Results:** We develop a deep learning method to correct non-uniform milling artifacts observed in FIB-SEM images. The proposed method is an image-to-image translation technique that can mitigate image distortions in an unsupervised manner. It conducts cross-plane learning within 3D image volumes without any ground truth annotations. We demonstrate the efficacy of our method on a real-world micro-wasp dataset, showcasing significantly improved image quality after correction with qualitative and quantitative analysis.

## Introduction

Focused Ion Beam Scanning Electron Microscopy (FIB-SEM) is an advanced Volume Electron Microscopy technology with remarkable isotropic 3D resolution [14, 27], leading to rapidly growing applications [10, 23, 4, 28]. Most Volume Electron Microscopes use diamond knives to section samples with a typical thickness of about 40 nm. In comparison, FIB-SEM uses a focused ion beam (FIB) to mill the sample surface, resulting in an effective “section” thickness of about 8 nm [3, 20], as shown in Fig. 1 (a). This state-of-the-art axial resolution is vital for accurately detecting, segmenting, and reconstructing fine structures of cells, such as organelles and synapses [10, 28].

**Fig. 1.**
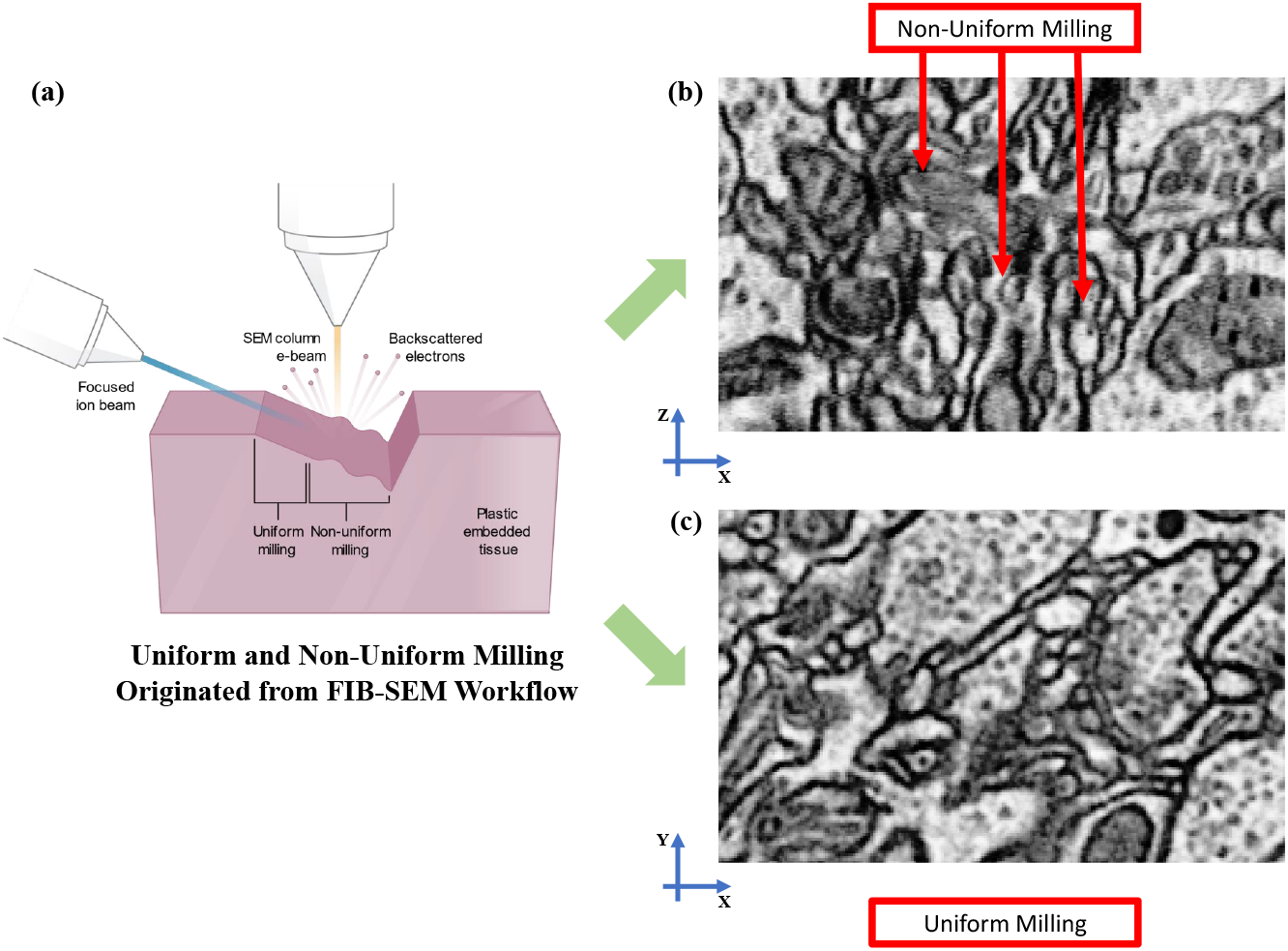
(a): Uniform and non-uniform milling originated from FIB-SEM workflow. The focused ion beam creates wave-like artifacts on the sample surface during the milling process. (b): An example (from X-Z plane) of non-uniform milling in a re-sectioned FIB-SEM image stack (sectioning plane is horizontal). Discontinuity between consecutively imaged sections is indicated by the red arrows. (c): An example (from X-Y plane) of uniform milling in a FIB-SEM raw image section stack.

### Non-Uniform Milling

Ideally, the milling thickness should be constant (uniform milling) across sample surfaces, and the block-face scanning should create naturally aligned images. In practice, however, both the physical properties, such as hardness, of the biological structures embedded in resin and the FIB milling force applied to the sample surface are uneven. One can think of the non-uniform milling process during FIB-SEM imaging as wind blowing through a lake’s surface, creating uneven waves (see Fig. 1 (a)). As a result, the milling thickness varies (non-uniform milling) across the sample surface, leading to image volumes with axial plane distortions [2, 12], as indicated in Fig. 1 (b). Such distortions may negatively impact the accurate reconstruction of small neuronal structures.

### Related Work

In [12], a fiducial marker system was introduced to measure slice geometry in FIB-SEM imaging. Researchers conclude that instrument and material instabilities may be the causes of the thickness variation. Such a method adds a layer of system complexity to the image analysis workflow. Researchers compared several classic methods for estimating the thickness of ultrathin tissue sections [7]. The included methods require additional physical measurements, making them inefficient and impractical for modern microscopy image analysis. [1] uses two software applications combined with a volume flattening procedure to deal with digitized volumes that have tears and thickness variations within tissue sections. [2] proposes to control the position of the ion beam during the milling process to reduce non-planar distortions. Another work [24] introduces an approach that uses image statistics alone, i.e., the standard deviation of the difference between pixel values as a function of distance, to help estimate the section thickness. [8] describes a traditional image processing method to estimate the milling thickness by computing the similarity or distance between neighboring sections. It assumes that the section thickness is uniform for each section. However, such an assumption does not hold for certain datasets, such as the micro-wasp dataset [4, 17] in this work, resulting in inferior performance. Despite these prior efforts, to the best of our knowledge, the problem of correcting non-uniform milling is still unsolved.

Benefitting from the boom of deep learning technologies [16], a great number of advanced image preprocessing methods were developed, such as image denoising [25], image restoration [26, 9], etc. Among these newly established techniques, image-to-image translation attracts our particular interest. It is a deep learning-based image processing method that transforms images from one domain to another, such as turning black-and-white photos into color or converting sketches into realistic images. In this field, two notable works are proposed in and [31], which develop generative adversarial networks (GANs) [6] to perform image translation in a supervised and unsupervised manner, respectively. Supervised image-to-image translation uses paired training data where each image in the source domain has a corresponding image in the target domain to learn the mapping, while unsupervised image-to-image translation uses unpaired training data to learn the mapping between domains without direct correspondences. Extensions of these methods are widely applied to various problems, e.g., biomedical data processing [29, 30, 18] and imaging process optimization [5, 26].

### Our Contributions

We propose an unsupervised cross-plane image-to-image translation pipeline to correct non-uniform milling without perfectly milled ground truth samples. Our algorithm is based on the following. Whereas non-uniform milling results in discontinuity in image re-sections in the axial direction (i.e., the X-Z and the Y-Z plane) of the 3D FIB-SEM volume, raw image sections (the X-Y plane) do not have such an issue (Fig. 1 (b) and (c)). Therefore, we formulate the problem as an unsupervised learning task to translate images from the X-Z/Y-Z domain (with artifacts) to the X-Y domain (without artifacts). Then, during the inference stage, given an image obtained from the X-Z/Y-Z plane, the model is able to reduce its line-like artifacts. The main contributions of this paper are as follows:

- We apply image-to-image translation methods to explicitly tackle the problem of non-uniform milling in FIB-SEM images in an end-to-end manner.
- The proposed method can mitigate image distortions in an unsupervised manner, i.e., it conducts cross-plane learning within 3D image volumes themselves without any ground truth image volumes.
- The performance of our method and a baseline method is evaluated qualitatively and quantitatively on a real-world micro-wasp FIB-SEM dataset. The results illustrate that the proposed method can significantly improve image quality and reduce the non-uniform milling artifact.

## Methods

In this section, we introduce an unsupervised cross-plane image-to-image translation method. The pipeline (Fig. 2) can be applied to correct image re-sections obtained along either the X-Z plane or the Y-Z plane. In the following contents, we only mention the Y-Z plane for the sake of clarity.

**Fig. 2.**
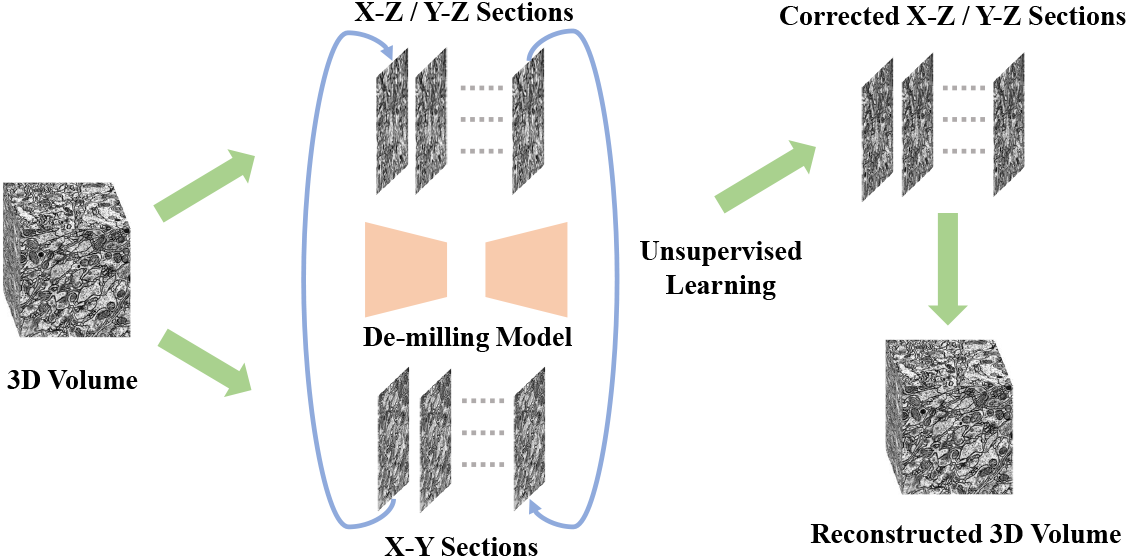
Unsupervised cross-plane image-to-image translation pipeline. A 3D FIB-SEM volume is sliced to create a bag of X-Y sections and a bag of X-Z/Y-Z sections. The X-Y sections are used as the training target of the de-milling model, while the X-Z/Y-Z sections are used as the input to the model. Through unsupervised cross-plane learning, the model produces corrected X-Z/Y-Z sections followed by reconstructing the sections into a 3D FIB-SEM volume.

### Unsupervised Cross-Plane Learning

After registration and contrast adjustment a serial stack of images acquired by FIB-SEM forms an image volume 𝒱. We decomposed the volume into two bags of 2D image sections along the X-Y and Y-Z planes. The volume 𝒱 can be expressed as a function 𝒱(*x, y, z*), where (*x, y, z*) represents the spatial coordinates within the volume. For each *z*, we obtain a 2D section in the X-Y plane, leading to a bag of 2D images denoted as *I*_*xy*_(*z*) = 𝒱(*x, y, z*). Similarly, for each *x*, we obtain sections in the Y-Z plane, leading to a bag *I*_*yz*_(*x*) = 𝒱(*x, y, z*).

Image sections in bag *I*_*yz*_(*x*) have the discontinuity issue due to non-uniform milling. To train a model that can correct such discontinuity, supervised image-to-image translation approaches often require access to one-to-one corresponding ground truth labels, i.e., perfectly milled images in our case. Unfortunately, these images cannot be obtained in practice.

To this end, we propose an unsupervised cross-plane learning approach. Specifically, we aim to train a de-milling model *M*_*θ*_ that learns the mappings between *I*_*yz*_(*x*) and *I*_*xy*_(*z*) in order to correct a Y-Z section to have a quality similar to an X-Y section, without the need to have one-to-one ground truth labels as the model is only learning the distribution of *I*_*xy*_(*z*).

During the inference stage, given *I*_*yz*_(*x*) from the test set, the trained model can output a bag of predictions *Î*_*yz*_ (*x*), whose non-uniform milling issue has been corrected, exhibiting similar quality to *I*_*xy*_(*z*). Then, we reconstruct 2D image sections from *Î*_*yz*_ (*x*) into a corrected 3D volume 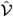 (*x, y, z*).

### De-Milling Model

Concretely, the objective of the de-milling model is to learn mapping functions between the Y-Z plane domain *A* and the X-Y plane domain *B* given unpaired training samples 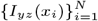 where *I*_*yz*_(*x*_*i*_) ∈ *A* and 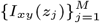 where *I*_*xy*_(*z*_*j*_) ∈ *B*. Following [31], the model includes two mappings (generators): 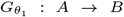, and 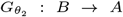, and two corresponding discriminators *D*_*A*_ and *D*_*B*_. The discriminator *D*_*A*_ aims to distinguish between real images *I*_*yz*_(*x*) and translated images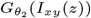, while *D*_*B*_ does the same for *I*_*xy*_(*z*) and 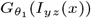.

### Adversarial Loss

To match the distribution of the generated images with the target domain, we apply adversarial losses [6] to both mapping functions. For the mapping function 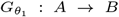 and its discriminator *D*_*B*_, the objective is expressed as:

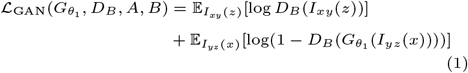

where 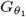 aims to fool *D*_*B*_ by generating images 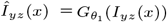 that have similar quality (i.e., reduced discontinuity) as *I*_*xy*_(*z*), while *D*_*B*_, on the other hand, tries to distinguish between generated images *Î*_*yz*_ (*x*) and real images *I*_*xy*_(*z*). Thus, we have an optimization goal as a min-max game: 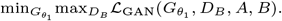.

Similarly, for the mapping of *B* → *A*, we have 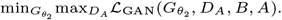.

### Cycle Consistency Loss

While adversarial losses ensure that the translated images look realistic, they do not guarantee that the learned mappings are meaningful or consistent. To address this, a cycle consistency loss is needed. This loss governs that translating an image from one domain to the other and back again should result in the original image. Mathematically, the forward (from *A* to *B* and back to *A*) and backward (from *B* to *A* and back to *B*) cycle consistency can be combined as the following loss:

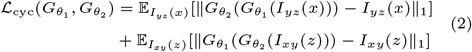

### Full Objective

The full objective for the de-milling model is a weighted sum of the adversarial losses and the cycle consistency losses:

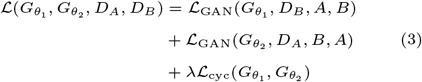

where *λ* controls the relative importance of the cycle consistency loss.

By minimizing this final loss, the de-milling model learns to correct the non-uniform milling artifacts in the Y-Z plane sections, producing high-quality images similar to the X-Y plane sections.

## Dataset

Micro-wasp (*Megaphragma vigianii*) is an interesting subject for neuroscience due to the small number of neurons in its brain, elaborate behavior, and miniaturization adaptations like the absence of nuclei in most neurons [19, 21]. With a special protocol, the whole head was prepared for electron microscopy imaging by staining with a heavy metal [22]. This sample was imaged using a customized FIB-SEM with a voxel size of 8 *×* 8 *×* 4 nm [27, 28]. The images were aligned using affine transformations, and the adjacent sections were combined by averaging, resulting in an isotropic voxel size of 8 *×* 8 *×* 8 nm. The images were preprocessed using HIFI-EM [15].

## Implementation Details

The de-milling model is trained for 100 epochs with a constant learning rate of 0.0002 and then for another 100 epochs to linearly decay the learning rate to 0. The optimizer is Adam [13], and the batch size is 1. All experiments are performed on an NVIDIA Tesla V100 GPU using the PyTorch deep learning library.

## Evaluation

We use the following methods to evaluate our proposed framework’s performance in correcting non-uniform milling in FIB-SEM images. The estimation of slice intervals between two sections for each location (*x, y*) is derived from the calculation of normalized cross-correlation (NCC) between patches centered on this location in the current and previous sections. Given two image sections, *P*_cur_(*x, y*) (the current section) and *P*_pre_(*x, y*) (the previous section) of size 135 *×* 135, the normalized cross-correlation NCC(*x, y*) is defined as:

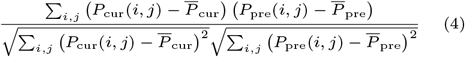

where *i, j* index the pixels in the patches, and 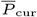 and 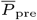 represent the mean intensities of the current and previous sections, respectively.

The normalized cross-correlation represents the similarity measurement between patches, with values closer to 1 indicating a higher similarity, and thus a shorter distance between the two sections at that location. To reduce high-frequency noise components and improve the quality of the NCC calculation, noise reduction is applied through a low-pass filter in the section plane.

The normalized cross-correlation values are then transformed into a distance metric using the arc-cosine function:

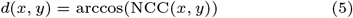

This transformation ensures that the resulting distances adhere to the triangular inequality, making them suitable for further analysis.

To interpret these distances throughout our image volume, and compare uncorrected and corrected variants using different methods, we want to map them into a normalized scale. We first calculate the mean value within a size-*n* one-dimensional window along the *z*-direction centered at each (*x, y, z*) location. It represents what is assumed to be a 1-unit interval and serves as a baseline for comparison:

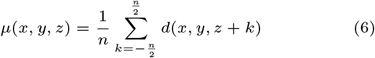

where *n* is the size of the window in the *z*-direction. Alternatively, a Gaussian-like filter, rather than the simple averaging filter shown here, could be used.

The normalized deviation of the slice interval from the average unit distance, for each (*x, y*) location, is estimated by dividing the distance metrics *d*(*x, y*) by the calculated mean value *µ*(*x, y, z*). The normalized slice interval *E*(*x, y, z*) is then given by:

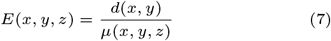

This normalization process provides a relative measure of the slice interval, indicating how much the interval at each (*x, y*) location deviates from the assumed unit interval.

## Experiments

In this section, we describe our main results. To demonstrate the efficacy of our proposed method, we applied the method from the closest prior work, i.e., Hanslovsky et al. [8], to our dataset and use the output as the baseline for comparison.

### Qualitative Results

Fig. 3 illustrates a comparative analysis between the original FIB-SEM input image, the corrected output from Hanslovsky et al. [8], and the result achieved by our proposed correction method. In both the X-Z and Y-Z planes, the input image reveals substantial non-uniform milling artifacts characterized by discontinuities and distortions that obscure finer cellular structures and complicate accurate segmentation. When applying the method by [8], some degree of artifact reduction is observed, particularly in the X-Z plane, where improvements in continuity and clarity can be noted. However, their approach leaves residual artifacts, especially visible in the Y-Z plane, where axial distortions continue to affect image fidelity. This limitation may stem from assumptions in their method regarding uniform section thickness, which do not hold across certain biological samples with variable milling properties.

**Fig. 3.**
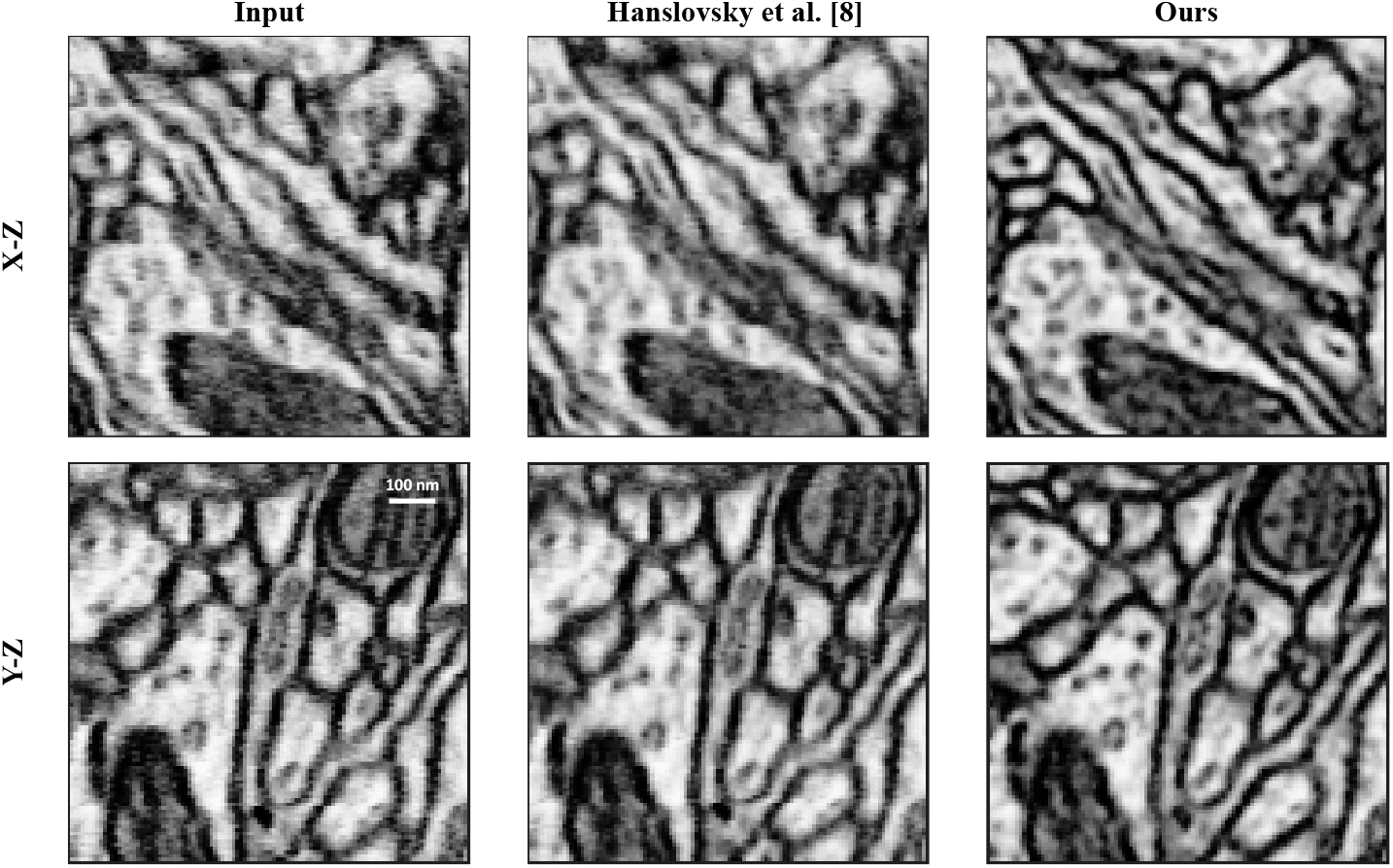
Comparison of non-uniform milling correction results for FIB-SEM images. Columns display (from left to right) the original input image, results using the method by Hanslovsky et al. [8] (baseline), and our proposed correction method. Rows show examples from different planes (X-Z and Y-Z). While the baseline approach reduces some artifacts, our method demonstrates a more significant improvement in pixel continuity and a stronger enhancement of the structural details of cellular membranes.

In contrast, our proposed approach employs an unsupervised cross-plane image-to-image translation technique that learns from 2D sections taken from the X-Y plane to better correct the non-uniform milling artifacts in both X-Z and Y-Z planes. Our method effectively restores continuity and removes distortions, allowing for a more faithful representation of cellular structures. Notably, compared to [8], the membrane regions benefit from sharper delineation and reduced noise, which are essential for subsequent analyses like organelle segmentation and structural tracing in applications such as connectomics [19]. Additionally, by minimizing the line-like artifacts caused by non-uniform milling, our method enhances the interpretability of biologically relevant features. This improvement is vital for high-resolution 3D reconstruction tasks, where accurate structural representation is paramount for understanding complex tissue architecture. It is clear that our approach significantly outperforms the baseline, making it a robust tool for large-scale, high-throughput microscopy data.

### Quantitative Results

To better understand the uniformity of milling thickness across different methods, we analyzed 2D sections of the 3D interval estimation maps, as shown in Fig. 4. These maps visualize the deviations in estimated slice intervals, providing a clear spatial representation of milling inconsistencies. The original interval estimation map reveals substantial deviations with pronounced irregularities distributed throughout the section. These variations highlight severe non-uniformities in milling thickness, with regions showing significantly higher or lower estimated intervals than the baseline value. The map generated by [8] shows some reduction in these inconsistencies, but high deviations are still obvious, demonstrating the limitations of the baseline method in achieving uniform milling. In contrast, our proposed X-Z and Y-Z corrections significantly reduce the overall deviation, as evidenced by more uniform interval estimation distributions across the map. Both corrections effectively address high-deviation regions, resulting in a smoother spatial representation. These results confirm that our approach successfully enhanced milling uniformity, outperforming the baseline method.

**Fig. 4.**
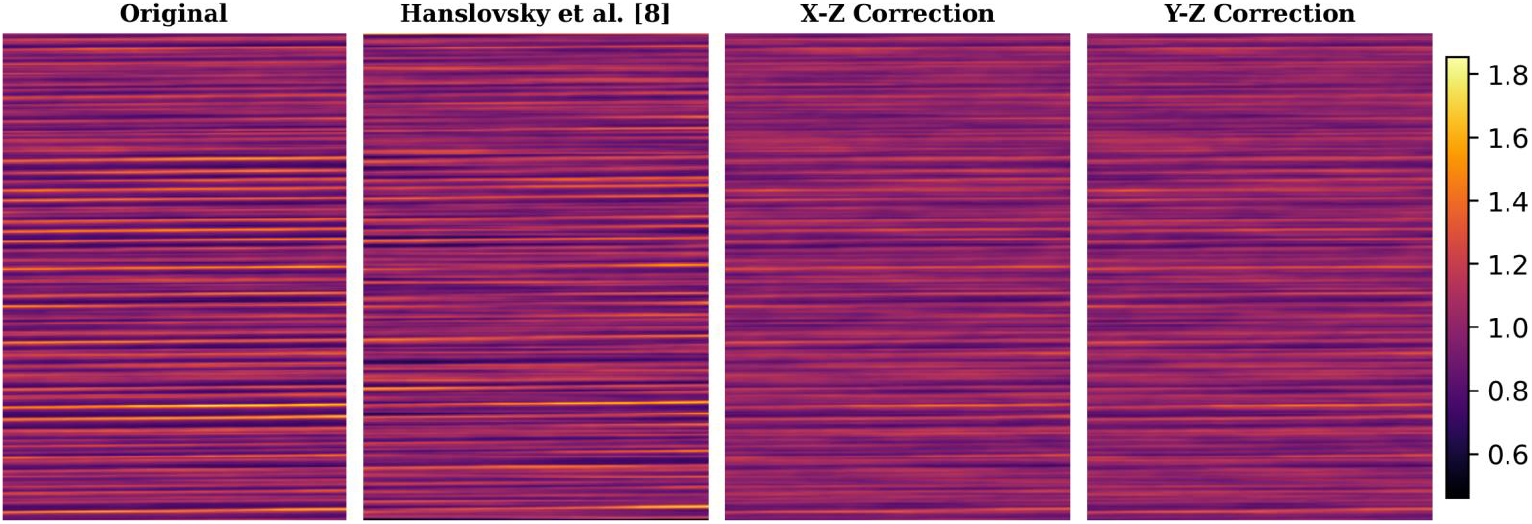
2D sections of the 3D interval estimation maps for different methods. The original map (left) shows substantial deviations and irregularities in estimated slice intervals, indicating severe non-uniformities. The baseline correction by [8] (second from left) reduces some deviations but leaves significant residual inconsistencies. The proposed X-Z correction (second from right) and Y-Z correction (right) demonstrate superior performance, achieving greater uniformity and effectively minimizing deviations across the section.

Furthermore, we calculated a probability density function (PDF), showing the normalized slice intervals and their deviation from a baseline unit distance (see Fig. 5). This metric provides a robust measure of how closely the corrected slice intervals align with an idealized uniform thickness, reflecting the reduction in milling artifacts. In the original dataset, the normalized slice intervals exhibited substantial deviations, indicating severe non-uniformities across the sample. After applying our proposed correction method, these variations were significantly reduced, resulting in a smoother and thinner distribution centered closer to the ideal scenario. This improvement demonstrates the capability of the method to enhance uniformity along the z-direction in FIB-SEM volumes. Notably, our method outperforms the baseline approach [8], which only reduces the deviation a little.

**Fig. 5.**
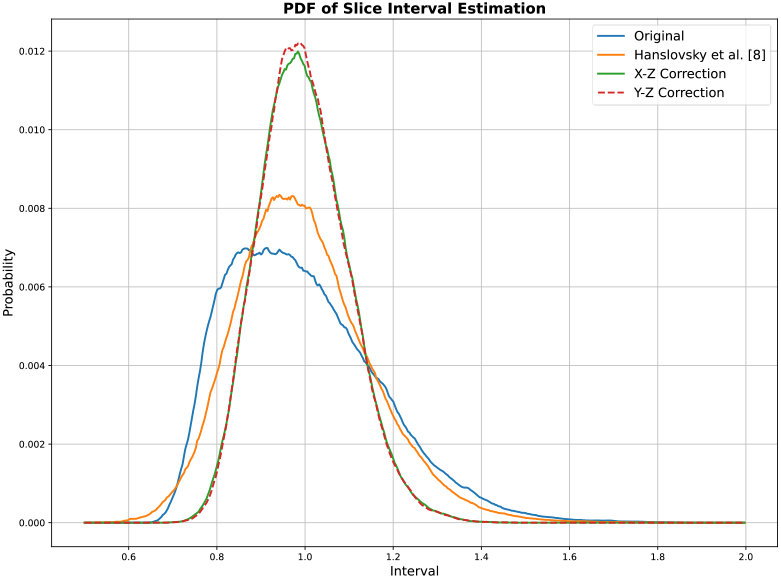
Quantitative evaluation of normalized slice intervals before and after correction. The original dataset exhibits significant deviations from uniformity, indicating non-uniform milling artifacts. The proposed method substantially reduces these deviations, resulting in a distribution closer to the idealized uniform thickness. In comparison, the baseline method by [8] shows only limited correcting ability, indicated by the tails of its PDF being fatter than those of our proposed method.

These quantitative results complement the qualitative improvements observed in the corrected images, further validating the effectiveness of the proposed approach in mitigating non-uniform milling distortions.

### Reconstruction Consistency

The robustness of the proposed method is further demonstrated by evaluating its ability to ensure reconstruction consistency in the X-Y plane. Fig. 6 shows a comparative analysis of the X-Y section from the reconstructed 3D volume using corrected X-Z/Y-Z sections against the ground truth X-Y section from the original 3D volume. In general, it can be observed that our method faithfully preserves structural details after the translation and the reconstruction process.

**Fig. 6.**
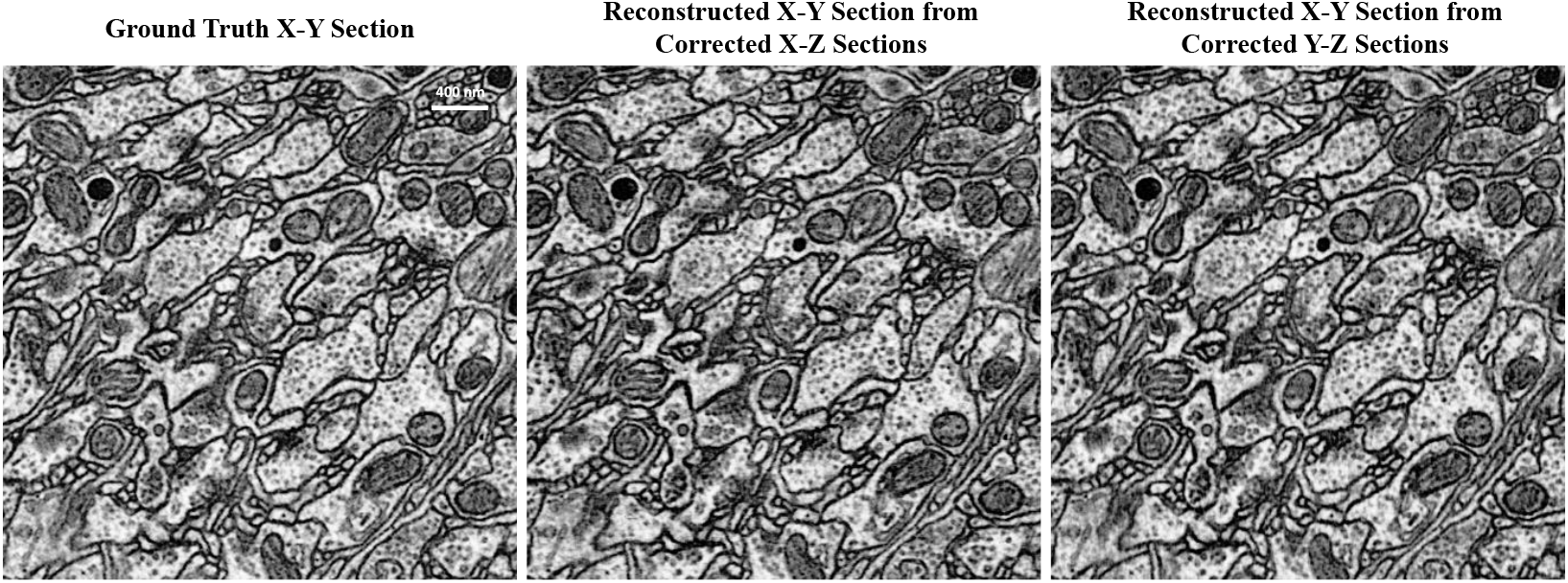
Evaluation of reconstruction consistency in the X-Y plane. The ground truth X-Y section (left) serves as a reference. The middle and right images are reconstructed X-Y sections using the corrected X-Z and Y-Z sections, respectively. Both reconstructions demonstrate high structural fidelity and alignment with the ground truth, preserving critical features such as membrane boundaries and intracellular details.

The ground truth X-Y section presents a clear and continuous representation of cellular structures. When the X-Y section is reconstructed from the corrected X-Z sections, the structural consistency closely aligns with the ground truth, with minor deviations observed in the finer membrane regions. Similarly, the reconstruction from the corrected Y-Z sections also demonstrates high fidelity, with no significant loss of information or introduction of artifacts. In both cases, the reconstructed X-Y sections maintain critical features such as membrane boundaries and intracellular details, which are vital for accurate 3D segmentation and biological interpretation.

The observed structural coherence between the reconstructed and ground truth X-Y sections highlights the method’s potential for application in high-resolution volumetric imaging workflows, where accurate reconstruction is essential for downstream analysis.

## Discussion

An observable effect of the proposed correction method is the enhanced visibility of cellular membranes, as illustrated in Fig. 7. Prior to correction, non-uniform milling artifacts distort the axial plane, leading to compromised continuity and structural fidelity in FIB-SEM images. After correction, however, the proposed unsupervised image-to-image translation technique mitigates these distortions, allowing for clearer delineation of fine structures, such as cellular membranes, that are critical for accurate neuronal segmentation. This improvement facilitates more reliable segmentation of organelles and cellular compartments, which is especially valuable in connectomics research.

**Fig. 7.**
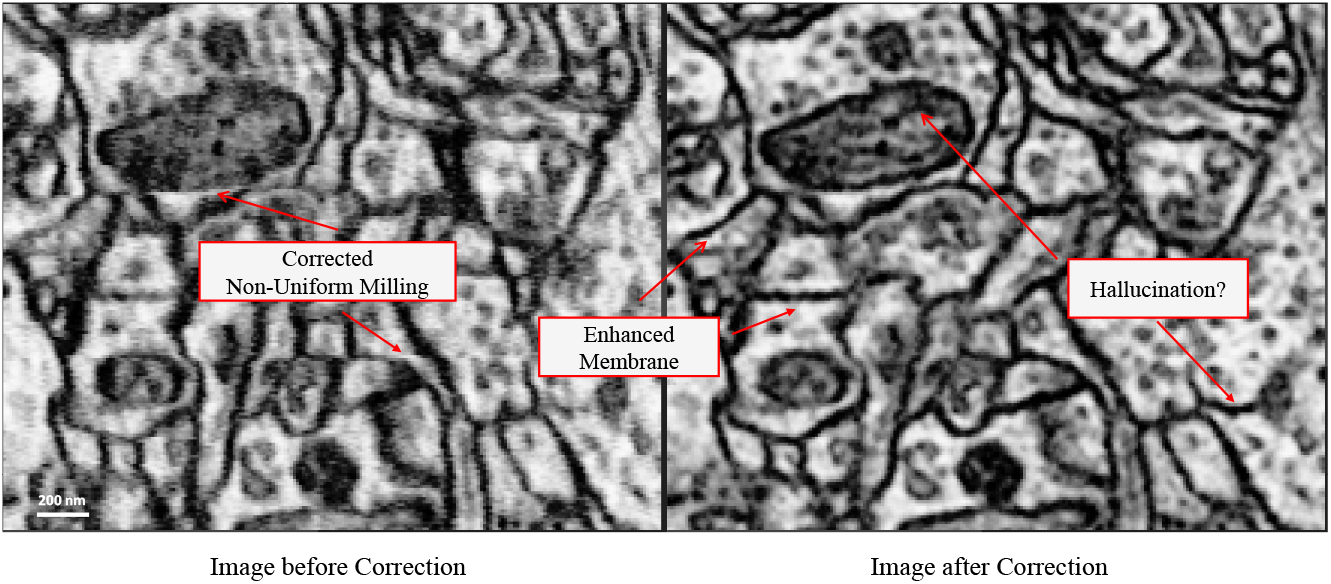
Comparison of FIB-SEM images before and after correction using the proposed method. Left: Original image with non-uniform milling artifacts that obscure membrane details and create discontinuities. Right: Corrected image, showing enhanced membrane clarity and structural continuity. The correction effectively mitigates milling-induced distortions, enabling clearer visualization of cellular structures. However, the possibility of artificial hallucination of features requires further validation to confirm the biological accuracy of the corrected image.

On the other hand, the observed enhancement in membrane clarity, while beneficial, also sometimes results in hallucinations, i.e., deep learning methods inadvertently generate features that might not be present in the original biological structures (Fig. 7). This phenomenon, commonly observed in generative adversarial models, underscores a critical need for verification: enhanced details in corrected images must be validated to ensure they represent true biological structures rather than artifacts introduced by the model. Rigorous validation techniques, possibly including comparison with ground-truth data or cross-referencing with alternative high-resolution imaging modalities, would strengthen confidence in the corrected images.

## Conclusion

In this paper, we present an unsupervised cross-plane image-to-image translation method to correct non-uniform milling artifacts in FIB-SEM image volumes. Our method demonstrates superior performance over previous approaches, as evidenced by enhanced structural fidelity in corrected images and significantly improved uniformity in slice intervals. By eliminating the need for ground-truth data, the proposed approach provides a scalable, efficient solution for high-throughput volumetric imaging workflows in connectomics.

## Author contributions statement

Y.L. and J.W. conceived the experiments. Y.L. and Y.K. conducted the experiments. Y.L., Y.K., S.H., E.S., and J.W. analyzed the results. All authors wrote and reviewed the manuscript. D.C., H.P., and J.W. supervised and supported the project.

## Acknowledgments

This project is supported by the Simons Foundation, Lingang Laboratory grant LGL-8998, NSF grant NCS-FO-2124179, and NIH grant 1U01NS132158.

